# Optimizing Network-Level TMS-fMRI: Benchmarking the TMS-Compatible Sushi MR Setup

**DOI:** 10.64898/2026.01.27.701271

**Authors:** Yiwu Xiong, Michael Burke, Lorena Melo, Kuri Takahashi, Maximilian Lueckel, Til O. Bergmann, Michael A. Nitsche, Erhan Genç, Emilio Chiappini

**Affiliations:** Leibniz Research Centre for Working Environment and Human Factors (IfADo), 44139, Dortmund, Germany; SensoriMotorLab, Jules-Gonin Eye Hospital - Fondation Asile des Aveugles, Department of Ophthalmology, University of Lausanne (UNIL), 1004, Lausanne, Switzerland; International Institute for Integrative Sleep Medicine (WPI-IIIS), Tsukuba Institute for Advanced Research (TIAR), University of Tsukuba, 305-8575, Tsukuba, Japan; Neuroimaging Center (NIC), Focus Program Translational Neuroscience (FTN), Johannes Gutenberg University Medical Center, 55131, Mainz, Germany; Leibniz Institute for Resilience Research (LIR), 55122, Mainz, Germany; Bielefeld University, University Hospital OWL, Protestant Hospital of Bethel Foundation, University Clinic of Psychiatry and Psychotherapy, 33617, Bielefeld, Germany; German Center for Mental Health (DZPG), Bochum, Germany

**Keywords:** Concurrent TMS-fMRI, MR receive coil, multi-echo fMRI, temporal SNR, network-level readout, interleaved TMS-fMRI

## Abstract

Concurrent TMS-fMRI can map how stimulation affects both the targeted cortex and connected brain-wide networks, but this requires MR receive hardware that allows TMS coil placement while preserving reliable whole-brain BOLD sensitivity. We developed and benchmarked a practical TMS-compatible “Sushi” MR receive setup assembled from two flexible 18-channel body arrays. Across six experiments, we tested functional readout validity, signal quality, and active TMS-fMRI compatibility. Resting-state fMRI (n = 12) and verbal N-back task-fMRI (n = 8) were acquired with Sushi, a commercially available 2×7-channel Surface setup, and a standard 64-channel head/neck array. Functional similarity to the 64-channel reference was quantified with spatial overlap, and multi-echo combination (MEcomb) was tested as a post-acquisition signal optimization strategy. Sushi recovered subject-specific resting-state networks that more closely matched the 64-channel reference than Surface, with no significant difference from the 64-channel test-retest reference. For task-fMRI, MEcomb increased task-map similarity for Sushi, whereas setup comparisons within each pipeline were not significant. MEcomb also improved resting-state similarity and increased temporal signal-to-noise ratio (tSNR) across receive setups. In phantom measurements, TMS coil placement produced spatially graded tSNR reductions relative to the no-TMS-coil condition, strongest near the coil. In one participant, active interleaved single-pulse TMS-fMRI over two cortical sites showed no detectable pulse-locked image artifacts; whole-brain MEcomb tSNR during active TMS-fMRI was reduced by 2.6-6.9% relative to the no-coil/no-stimulation reference. Together, Sushi and MEcomb provide complementary hardware and processing tools for TMS-compatible whole-brain fMRI. This combination supports network-level functional readouts while preserving feasibility for active interleaved TMS-fMRI.

## 1 INTRODUCTION

Transcranial magnetic stimulation (TMS) perturbs a focal cortical site embedded in distributed cortical and subcortical networks, inducing effects that can propagate locally and across connected nodes (Bergmann & Hartwigsen, 2021; Ferbert et al., 1992; Fox et al., 1997; Momi et al., 2021; Tik et al., 2023; Valero-Cabré et al., 2017; Xia et al., 2024). Exploring and validating these local and downstream effects requires a readout that is not restricted to the stimulation target but provides sufficiently reliable sensitivity across distributed brain regions. Concurrent TMS and functional magnetic resonance imaging (fMRI) promises such a readout by combining focal perturbation with whole-brain BOLD imaging, allowing local and distal effects of TMS to be mapped during stimulation (Bergmann et al., 2021; Bestmann et al., 2008; Bohning et al., 1998).

The quality of the MR receive hardware is therefore central to the inferences that can be drawn from concurrent TMS-fMRI. Reliable interpretation requires sufficient BOLD sensitivity not only at the stimulated site, but also across the distributed networks through which TMS effects may propagate (Bergmann et al., 2021; Woolgar et al., 2025). In practice, this promise remains only partially fulfilled because the MR-receive design imposes a critical trade-off between homogeneous whole-brain BOLD sensitivity and accessibility and flexibility of TMS coil placement. Standard multi-channel head coils provide high SNR and support accelerated fMRI through parallel imaging and multiband acquisition, but their close-fitting geometry leaves little or no room for the TMS coil (Navarro de Lara et al., 2015). TMS-compatible solutions, in contrast, provide access to the scalp yet often sacrifice signal quality, spatial homogeneity, or parallel imaging performance (Bergmann et al., 2021; Jackson et al., 2024; Woolgar et al., 2025). Filling this hardware gap is essential for concurrent TMS-fMRI (Bergmann et al., 2021; Mizutani-Tiebel et al., 2022; Schuler & Hartwigsen, 2025; Woolgar et al., 2025).

The main currently available TMS-compatible MR receive options include single-channel birdcage coil and multi-channel surface-array solutions. Birdcage coils provide broad and relatively homogeneous coverage, but with low SNR and no support for parallel-imaging acceleration. A popular alternative in TMS-fMRI research is the 2×7-channel TMS-specific surface-array setup (hereafter “Surface”; Navarro de Lara et al., 2015). This setup combines one small receive array positioned between the TMS coil and the scalp with a second array placed elsewhere on the head to extend coverage. Its multi-channel design supports accelerated imaging, but sensitivity remains concentrated near the receive arrays, leading to reduced SNR in more distant regions and pronounced spatial inhomogeneity (Jackson et al., 2024; Riddle et al., 2022; Woolgar et al., 2025).

Emerging multi-element hardware concepts have started to address these limitations (see Woolgar et al., 2025 for a broader overview). These include repurposed or customized flexible arrays that wrap around the head (Assem et al., 2025; Tik et al., 2025), RF-cap designs in which receive elements are attached to a tight-fitting cap (Navarro De Lara et al., 2025), and integrated TMS-MR receiver concepts in which multiple stimulation elements are embedded within the receive-coil housing to allow electronic steering of the stimulation focus (Navarro de Lara et al., 2023; Souza et al., 2025). These approaches show that the field is moving toward higher-channel, more flexible, and more whole-brain-oriented solutions for concurrent TMS-fMRI. However, a key validation gap remains: many new setups have been assessed mainly with technical image-quality metrics, whereas fewer have been tested against a high-quality reference coil using subject-level functional network readouts. In addition, several solutions remain prototypes or are not yet widely deployable across MR sites.

Here, we developed and benchmarked a practical low-cost setup for concurrent TMS-fMRI aimed at whole-brain network imaging and tested it together with post-acquisition multi-echo optimization (Posse et al., 1999) to improve functional readout quality. The Sushi setup combines two 18-channel flexible body arrays mounted around the head, leaving space for TMS coil placement while preserving multi-channel receive coverage and multiband imaging capability. Because these arrays are available at many MR sites as standard scanner equipment, the setup is intended to be readily deployable with minimal additional hardware. We benchmarked Sushi against the Surface setup and the standard non-TMS-compatible Siemens 64-channel head/neck array. Our main emphasis was functional readout validity for network neuroscience. We therefore tested whether a TMS-compatible setup can yield subject-specific functional maps similar to those obtained with a high-quality standard receive coil.

Across six experiments, the Sushi setup was validated at three levels: functional readout validity, signal quality, and active TMS-fMRI compatibility.

We first benchmarked Sushi against the Surface setup and the 64-channel reference using functional readouts. Experiment 1 examined whether each setup recovered eight subject-specific canonical resting-state networks (RSNs; Smith et al., 2009), providing network-level validation independent of a specific task. Experiment 2 examined whether the same setups preserved task-evoked BOLD patterns during a verbal N-back paradigm expected to recruit a distributed frontoparietal working-memory network (Emch et al., 2019; Rottschy et al., 2012). In both experiments, within-subject similarity to the 64-channel reference was quantified using spatial overlap, indexed by the Dice-Sørensen coefficient (DSC), with lower DSC indicating reduced correspondence with the benchmark maps (Frankford et al., 2021). Two processing strategies aimed at improving BOLD readouts under spatially inhomogeneous receive sensitivity were also evaluated. First, an intensity-homogenization procedure was applied to reduce sensitivity-profile bias and improve anatomical alignment. Second, multi-echo combination (MEcomb) was used to improve temporal stability and BOLD sensitivity in multi-echo EPI data (Posse et al., 1999). In Experiment 3, the signal-quality basis of the functional results was examined by benchmarking tSNR in human data across setups.

We then tested the Sushi setup under conditions directly relevant for concurrent TMS-fMRI. Because concurrent TMS-fMRI necessarily places TMS hardware close to the imaged object, Experiment 4 quantified the practical tSNR cost of TMS coil placement in controlled phantom measurements using the same benchmark sequence. Experiment 5 directly tested active concurrent TMS-fMRI compatibility: pulse-related image artifacts were assessed in a human participant using an interleaved sequence optimized for Sushi performance, and tSNR was quantified during active TMS relative to a no-coil/no-stimulation reference. Experiment 6 characterized the tSNR of this interleaved sequence in the phantom with and without the TMS coil.

Together, this design tested whether a readily deployable TMS-compatible receive setup, combined with MEcomb, can support network-level functional readouts while remaining compatible with active interleaved TMS-fMRI.

## 2 METHODS

### 2.1 Participants

Twelve healthy right-handed adults (6 female; mean ± SD age = 29.4 ± 5.1 years) participated in Experiments 1 and 3 and completed the resting-state fMRI sessions. Eight of these participants also completed Experiment 2, which involved task-fMRI sessions (4 female; mean ± SD age = 29.2 ± 4.9 years). One of these participants (male; age = 40 years) also took part in Experiment 5, which involved active concurrent TMS-fMRI. None of the participants had contraindications to MRI; the participant of Experiment 5 had no contraindications to TMS either. Participants had no relevant medical conditions and were not taking ongoing medication. All procedures were approved by the local ethics committee at IfADo (prot. 232/2023) and were conducted in accordance with the Declaration of Helsinki. All participants provided written informed consent and received financial compensation.

### 2.2 Materials and procedures

Six experiments were conducted to test the performance of the Sushi setup. Experiments 1 and 2 formed the matched human functional benchmark, in which the same benchmark fMRI sequence was used with the 64ch, Surface, and Sushi setups. Experiment 3 used the resting-state data from Experiment 1 to quantify whole-brain and RSN-level tSNR across receive setups. Experiment 4 used the same benchmark sequence in a phantom with the Sushi setup to quantify tSNR and the passive effect of TMS coil placement. Experiment 5 tested the feasibility of active concurrent spTMS-fMRI in one participant using an interleaved sequence by assessing TMS-induced artifacts in the EPI data and by quantifying tSNR under active stimulation conditions. Experiment 6 used the same interleaved sequence in the phantom to quantify tSNR and the passive effect of TMS coil placement without pulse delivery.

All fMRI experiments used multi-echo EPI acquisitions. To assess the contribution of MEcomb, we compared the optimally combined time series with the medium-TE time series, which corresponds most closely to the conventional BOLD-sensitive time of echo (TE) range used in single-echo fMRI.

The fMRI scans were performed on a 3T Magnetom Prisma scanner (Siemens Healthineers, Erlangen, Germany) at IfADo. When TMS hardware was present, the MR environment was configured to minimize electromagnetic interference. An MRI-B91 figure-of-eight TMS coil (MagVenture, Farum, Denmark) connected to a MagPro X100 stimulator (MagVenture) with dynamic leakage current suppression was used, together with a filter box containing low-pass filters (<100 kHz) addressing known artifact sources related to residual leakage currents and RF noise carried through the TMS system (Bungert et al., 2012; Weiskopf et al., 2009).

#### 2.2.1 Experiments 1-3: human benchmark

Three MR receive setups were tested in the human benchmark experiments: (i) the standard Siemens 64-channel head/neck receive array (64ch; Fig. 1A); (ii) the Surface setup with two TMS-dedicated 7-channel surface arrays (MagVenture), one positioned between the TMS coil and the scalp and the other on the opposite side of the head to extend coverage (Fig. 1B); and (iii) the custom Sushi setup with two 18-channel body arrays (Siemens Healthineers) mounted around the head in a custom holder (Fig. 1C-D). Each participant completed four sessions: two with the 64ch setup (test-retest reference; 64ch-1, 64ch-2), one with the Surface setup, and one with the Sushi setup. The 64ch-1 session was always acquired first, and the order of the remaining sessions was randomized.

**Figure 1.**
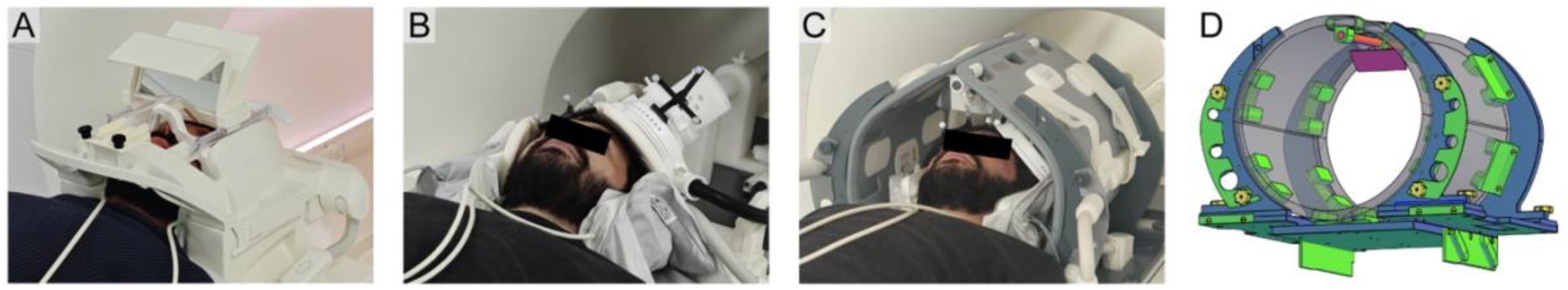
MR receive setups. (A) Standard 64-channel head/neck receive array. (B) Surface setup used in the study, with two 7-channel surface arrays and the TMS coil in place. (C) Sushi setup, with two flexible 18-channel body arrays mounted in the custom holder and the TMS coil in place. (D) Rendering of the Sushi holder with color-coded components: grey, flexible 18-channel body arrays; blue, cut plastic plates (PVC); green, 3D-printed parts (PLA); yellow, nylon screws; purple and orange, mirror and mirror support.

In the TMS-accessible setups, the TMS coil was placed tangentially over a left central scalp region approximating the primary motor cortex (M1) hand area in half of the participants (six in rs-fMRI, four in task-fMRI), and over a left lateral prefrontal scalp region approximating the dorsolateral prefrontal cortex (dlPFC) in the other half. The coil was fixed with an adjustable holder (MagVenture). Inflatable cushions and a vacuum head pillow were used to stabilize the head and, in Surface setup sessions, the second surface array. No TMS pulses were delivered for these experiments.

In Experiment 1, one 10-min rs-fMRI run was acquired for each setup. Participants rested in a darkened room with eyes closed and were instructed to remain still and awake, without engaging in any specific mental activity.

In Experiment 2, participants performed a verbal N-back task (1-back, 3-back, and 4-back) in a block design. Each participant completed two runs lasting approximately 7.5 min each. Each run included two blocks of each N-back condition, resulting in four blocks per condition across runs. The 3-back condition was included as an additional condition but was not used in the present analyses. Analyses focused on the 4-back > 1-back contrast. Each task block was preceded by a 12-s fixation period and a 7-s instruction screen. The task block lasted 54 s and comprised 30 trials (300-ms letter presentation followed by a 1500-ms fixation/response period), including 6 target and 24 non-target trials presented in pseudorandom order. Participants responded to target and non-target trials with the right index and middle fingers using a multi-button response box (Psychology Software Tools, Pittsburgh, USA); such response mapping was counterbalanced across participants. Stimuli were presented on a screen (MedRes GmbH, Cologne, Germany) at the back of the bore and viewed via a mirror. The task was implemented in MATLAB (R2021b; The MathWorks Inc., Natick, USA) using Psychtoolbox (PTB3) and synchronized with the MR trigger.

Experiment 3 used the rs-fMRI data from Experiment 1 to quantify whole-brain and RSN-level tSNR. No additional data were acquired for this experiment.

#### 2.2.2 Experiment 4: phantom tSNR with the benchmark sequence

Experiment 4 used the same benchmark sequence as Experiments 1-3, but data were acquired in a cylindrical Siemens phantom (1.9-L plastic bottle Siemens phantom filled with Solution N) using the Sushi setup. The purpose was to obtain an objective and reproducible tSNR reference under controlled conditions and to quantify the passive effect of TMS coil placement on tSNR. Runs were acquired with and without the TMS coil positioned against the phantom. Vitamin-E markers visible on the T1-weighted image were placed in the TMS coil to identify the exact coil position. Again, no TMS pulses were delivered for this experiment.

#### 2.2.3 Experiment 5: active concurrent spTMS-fMRI with interleaved sequence

In Experiment 5, the Sushi setup was tested for active concurrent single-pulse TMS-fMRI (spTMS-fMRI) compatibility. Single TMS pulses were delivered between slice packets during fMRI acquisition. To allow pulse interleaving, an interleaved EPI protocol optimized for the Sushi setup was implemented, with improved spatial resolution while maintaining a fast TR and balanced tSNR. Two 5-min runs were acquired, one targeting left M1 and one targeting left dlPFC, with 50 spTMS pulses per run. The M1 location was defined functionally as the site at which spTMS elicited the largest motor evoked potential in the first dorsal interosseous muscle. The dlPFC location was defined according to the MNI coordinates x = -38, y = 44, z = 26 (Blumberger et al., 2018) and localized using neuronavigation (Localite GmbH, Bonn, Germany). TMS intensity was deliberately set at 85% of maximum stimulator output for both targets; stimulation was well tolerated by the participant. An additional no-coil/no-stimulation reference run was acquired with the same interleaved sequence for descriptive tSNR comparisons. Therefore, tSNR reductions relative to the reference run were interpreted as reductions under active TMS-fMRI acquisition conditions rather than as passive coil-placement effects. The exact TMS coil position was indicated by vitamin-E markers.

#### 2.2.4 Experiment 6: phantom tSNR with interleaved sequence

Experiment 6 used the interleaved sequence from Experiment 5 in the same cylindrical Siemens phantom. Data were acquired with the Sushi setup, with and without the TMS coil positioned against the phantom. The exact TMS coil position was indicated by vitamin-E markers. Here, no TMS pulses were delivered. The purpose was to quantify controlled phantom tSNR for the interleaved sequence and to estimate the passive tSNR cost associated with TMS coil placement independently of pulse delivery.

### 2.3 MRI acquisition

High-resolution T1-weighted images were acquired with an MPRAGE sequence (TR = 2530 ms, TE = 2.36 ms, flip angle = 7°, FoV = 256 mm, voxel size = 1 mm³).

Experiments 1-4 used the benchmark EPI protocol. T2*-weighted EPI time series were acquired with a multiband-accelerated multi-echo sequence (Xu et al., 2013). The sequence sampled three TEs = 13, 35, and 56 ms and had TR = 1250 ms, flip angle = 48°, multiband factor = 3, in-plane FoV = 215 mm, voxel size = 3.2 × 3.2 × 3.5 mm, and AP phase encoding.

Experiments 5 and 6 used an interleaved EPI protocol optimized for concurrent spTMS-fMRI with the Sushi setup. T2*-weighted EPI time series were acquired using an accelerated multi-echo sequence with TR = 2000 ms, TEs = 11, 31, and 51 ms, flip angle = 80°, multiband factor = 3, in-plane FoV = 220 mm, voxel size = 2.2 × 2.2 × 2.4 mm, and AP phase encoding. The volume acquisition time was 1436 ms, leaving 564 ms of acquisition gap within each TR. This gap was evenly distributed across 18 multiband packets, allowing spTMS delivery immediately after the end of one packet and approximately 30 ms before the onset of the next packet.

For both EPI protocols, three additional spin-echo EPI volumes with reversed phase encoding (PA) were acquired to enable distortion correction. In Experiments 1-2, two 2D-GRE reference images (TR = 767 ms, TE = 4.92 ms, flip angle = 60°) were acquired for intensity homogenization: one using the integrated scanner body coil as receiver and one using the corresponding receive setup.

### 2.4 MRI data processing

#### 2.4.1 Multi-echo combination and image intensity homogenization

For MEcomb, the echo-specific time series were optimally combined into a single time series using the TE-dependent signal model of multi-echo EPI (Posse et al., 1999). Echo combination was implemented using the multi-echo combining functions of tedana (DuPre et al., 2021), without subsequent ICA-based component classification or denoising.

For Experiments 1-3, both the medium-TE and MEcomb time series were intensity-homogenized before registration and downstream analyses. This yielded two preprocessing pipelines: a homogenized medium-TE pipeline and a homogenized MEcomb pipeline. For Experiments 4-6, medium-TE values were retained as the reference for descriptive tSNR checks, while MEcomb was used as the primary output.

Intensity homogenization was applied to both preprocessing pipelines of Experiments 1-3 to reduce receive-sensitivity bias and support robust EPI-to-T1 registration. The correction field was computed from two GRE reference images acquired with the integrated body coil and the corresponding receive coil. Following (Narayana et al., 1988), both images were Gaussian-smoothed (σ = 5 voxels), and their ratio was used to estimate a smooth, time-invariant receive-sensitivity correction field. This field was aligned to the EPI data and applied to each volume. Because the correction factor was constant over time, temporal structure was preserved, whereas voxel-wise signal amplitudes were rescaled. This procedure was designed to reduce spatial intensity bias, especially in the Surface setup, and to support robust anatomical alignment (Fig. S1).

#### 2.4.2 Preprocessing workflow

Preprocessing was implemented in MATLAB (v. R2021a) and Python (v. 3), calling FSL (v. 6.0.7; Jenkinson et al., 2012), tedana (DuPre et al., 2021), ANTs (v. 2.3.5; Avants et al., 2011). For each EPI dataset, the same preprocessing workflow was used across receive setups where applicable. For human data, the temporal mean EPI was computed, a brain mask was derived, and the mask was applied to all volumes to remove non-brain tissue. Motion and susceptibility-distortion corrections were estimated and applied in native EPI space using the AP time series as functional input and the opposite phase-encoding PA images as reference. Corrected human data were coregistered to the corresponding T1-weighted image and transformed to MNI space for group-level and template-based analyses. Further technical details are reported in the Supplementary Materials.

#### 2.4.3 Functional and tSNR processing

For Experiment 1, rs-fMRI, single-subject ICA was performed with FSL MELODIC. IC spatial maps were registered to MNI space and compared with the eight selected Smith et al. (2009) RSN templates. These included the visuo-medial network (VisMN), visuo-lateral network (VisLN), default mode network (DMN), sensorimotor network (SMN), auditory network (AudN), executive control network (ECN), and right and left frontoparietal networks (FPN-r, FPN-l). The occipital pole visual and cerebellar networks were excluded to avoid overrepresentation of visual systems and because of incomplete cerebellar coverage in standard fMRI acquisitions. ICs with spatial correlation r < 0.10 to all selected templates were discarded. The remaining ICs were visually inspected, and obvious noise components were removed only when independently classified as noise by all three inspectors (E.C., Y.X., M.B.). The retained IC maps were entered into a spatial GLM against the Smith et al. templates. For each RSN template, the IC with the largest beta coefficient was selected as the corresponding subject-specific RSN map. Selected maps were thresholded at Z > 3 and binarized before DSC computation.

For Experiment 2, task-fMRI activation maps were derived from corrected, normalized, and smoothed EPI data using a block-design GLM in SPM12. Runs were modeled as separate sessions. Regressors for 1-back, 3-back, and 4-back blocks were convolved with the canonical HRF, and a 108 s high-pass filter was applied. The contrast of interest was 4-back > 1-back, yielding one first-level t-statistic map for each subject, setup, and preprocessing pipeline. These maps were used for DSC similarity analyses against the 64ch-1 reference. Group-level activation maps were also computed for qualitative display and to verify that DSC values reflected overlap within the expected working-memory network (Emch et al., 2019; Rottschy et al., 2012).

For Experiment 3, voxelwise tSNR maps were computed from the preprocessed rs-fMRI time series acquired in Experiment 1 as the temporal mean divided by the temporal standard deviation (Welvaert & Rosseel, 2013). Human tSNR maps were transformed to MNI space, brain-masked, and summarized as whole-brain means and as means within canonical RSN masks. RSN masks were obtained from the Smith et al. (2009) RSN templates thresholded at Z > 3 and corresponded to the same eight networks used for the RSN benchmark: VisMN, VisLN, DMN, SMN, AudN, ECN, FPN-r, and FPN-l.

For Experiments 4 and 6, phantom tSNR maps were analyzed in native EPI space. Phantom masks were derived from the thresholded mean EPI image. Mean local tSNR was extracted from two spherical ROIs with 25-mm radius positioned along the projected TMS coil axis as indicated by the vitamin-E markers placed in the TMS coil. The superficial ROI was centered 25 mm from the coil-facing phantom surface, and the deeper ROI was centered 75 mm from the same surface, so that the ROIs did not overlap. The same whole-phantom masks and local ROIs were applied to the corresponding no-coil and with-coil runs. Because of the different voxel sizes of the benchmark and interleaved sequences, the full spherical ROIs contained approximately 1850 voxels in Experiment 4 and 5580 voxels in Experiment 6.

For Experiment 5, pulse-related image artifacts were assessed in the MEcomb data using a slice-wise image-stability metric. For each axial slice, peak signal-to-noise ratio (PSNR) was computed across consecutive volume transitions, with lower PSNR indicating larger image changes. Slices were grouped into multiband packages according to the known MB = 3 acquisition scheme. A package was flagged when PSNR fluctuations were highly correlated across package members, and slice-level events were identified when PSNR showed a sudden drop across adjacent transitions. Detected events were evaluated relative to the recorded TMS pulse timing. tSNR was computed from these active spTMS-fMRI runs and from an additional no-coil/no-stimulation reference run. These data were coregistered and analyzed in 2-mm MNI space. The M1 and dlPFC TMS coil centers were derived from the vitamin-E markers. As in Experiments 4 and 6, local tSNR was extracted from superficial and deeper spherical ROIs with 25-mm radius positioned below the projected coil center for each target. The ROIs contained approximately 8050 voxels. Given the presence of non-brain tissue between the TMS coil and the nearest cortical tissue, the superficial and deeper ROI centers were located approximately 50 and 100 mm from the TMS coil surface for the M1 target, and 45 and 95 mm for the dlPFC target. No BOLD responses to the TMS pulses were modeled for this experiment.

Additional details on analysis and processing are provided in the Supplementary Materials.

### 2.5 Data analysis

To compare functional readouts across setups in Experiments 1 and 2, spatial-overlap analyses were performed using the 64ch-1 session as the within-subject reference. DSC was therefore computed between each comparison setup and 64ch-1. The comparison setup factor included 64ch-2, Sushi, and Surface. The 64ch-2 versus 64ch-1 comparison served as the test-retest reference for the 64-channel setup and therefore indexed expected stability of the functional readout. No participants were excluded from any completed experiment.

For Experiment 1, subject-specific RSN maps were thresholded at Z > 3 and binarized before DSC computation. This threshold was chosen to reduce obvious noise while preserving single-subject network overlap. DSC values were computed between each comparison setup and the 64ch-1 reference, separately for each subject, RSN, and preprocessing pipeline. RSN similarity was then analyzed across setups with DSC as the dependent variable. The model included SETUP, PIPELINE, RSN, and their interactions as fixed effects, with subject as a random intercept: *DSC ∼ SETUP × PIPELINE × RSN + (1 | SUBJ)*. SETUP included 64ch-2, Sushi, and Surface. PIPELINE included the medium-TE and MEcomb pipelines. RSN included the eight canonical networks used for the benchmark. More complex random-effects structures did not yield stable convergence or improved model fit, so the random-effects structure was kept as intercept-only.

For Experiment 2, first-level maps of the 4-back > 1-back contrast were converted to Z maps, thresholded at Z > 3, binarized, and compared with the corresponding 64ch-1 reference map within subject. Task-activation similarity was then analyzed across setups with DSC as the dependent variable. The model included SETUP, PIPELINE, and their interaction as fixed effects, with subject-specific SETUP slopes: *DSC ∼ SETUP × PIPELINE + (SETUP | SUBJ)*. Adding subject-specific SETUP slopes improved model fit relative to an intercept-only structure (ΔAIC = -57.1; χ²(5) = 67.1, p < 0.001), so this random-effects specification was retained.

For Experiment 3, whole-brain tSNR analyses used all four setup sessions: 64ch-1, 64ch-2, Sushi, and Surface. In contrast to the DSC analyses, 64ch-1 was not used as a reference here but entered the model as one setup level. Whole-brain tSNR was analyzed with SETUP, PIPELINE, and their interaction as fixed effects, with subject-specific SETUP slopes: *tSNR ∼ SETUP × PIPELINE + (SETUP | SUBJ);* adding subject-specific SETUP slopes improved model fit compared with an intercept-only model and was therefore retained. RSN-specific tSNR was analyzed with the same structure, but with RSN added as a fixed factor: *tSNR ∼ SETUP × PIPELINE × RSN + (SETUP | SUBJ)*.

For Experiments 4-6, tSNR was summarized descriptively because these experiments consisted of phantom measurements or single-participant feasibility data. We reported mean tSNR and the percentage tSNR reduction relative to the no-coil reference, computed as *100 × (1 − tSNRcoil/tSNRno-coil)*, for the whole brain or phantom volume and for the local ROIs.

For Experiment 5, pulse-related artifact analyses were summarized descriptively. A multiband slice package was flagged when the maximum pairwise correlation between PSNR time courses of package members exceeded 0.90. Within flagged packages, a slice-level event was defined as a PSNR drop greater than 1.0 dB across adjacent transitions.

Inferential analyses were conducted in R (v. 4.5.1; R Core Team, 2021) using lme4, lmerTest, and emmeans. Outcomes were analyzed with linear mixed-effects models. Fixed effects were tested with Type-III tests using Satterthwaite-approximated degrees of freedom. Model assumptions were evaluated by visual inspection of residual plots. Unless otherwise specified, post hoc contrasts are reported as differences in estimated marginal means (Δ) with standard errors (±SE). The alpha threshold was set at 0.05, and post-hoc p-values were Holm-corrected for multiple comparisons. Effect sizes for fixed effects are reported as partial eta squared (ηp²), computed from the F tests.

## 3 RESULTS

### 3.1 Experiment 1

As a descriptive check, RSN maps from the 64ch-1 reference showed consistent correspondence with the RSN templates (mean r = 0.43 ± 0.08 for medium-TE; r = 0.44 ± 0.1 for MEcomb), supporting its use as a functional reference for the DSC computations (see Table S1 and Fig. S2).

The LMM on DSC revealed significant main effects of SETUP (F_2,564_ = 47.1, p < 0.001, ηp^2^ = 0.14), PIPELINE (F_1,564_ = 7.0, p = 0.008, ηp^2^ = 0.01), and RSN (F_7,564_ = 6.7, p < 0.001, ηp^2^ = 0.08). A significant SETUP × RSN interaction was also present (F_14,564_ = 4.0, p < 0.001, ηp^2^ = 0.09). Other effects were not significant (p ≥ 0.28). The 64ch-2 DSC indexes test-retest similarity between 64ch-2 and 64ch-1, providing a within-setup reference for comparing Sushi and Surface similarity to the 64ch benchmark coil. MEcomb significantly increased similarity (DSC Δ = 0.04 ± 0.01, p = 0.012; Fig. 2A). In post hoc comparisons, Sushi did not significantly differ from 64ch-2 (Δ = 0.03 ± 0.02, p = 0.095), whereas the Surface setup was significantly lower than both 64ch-2 (Δ = 0.16 ± 0.02, p < 0.001) and Sushi (Δ = 0.13 ± 0.02, p < 0.001; Fig. 2B). More interestingly, post hoc comparisons of the SETUP × RSN interaction showed that the Surface setup performed significantly worse than the other setups in the auditory, DMN, and visual networks (p ≤ 0.001). Sushi did not significantly differ from 64ch-2 in any RSN (p ≥ 0.08; Fig. 2C).

**Figure 2.**
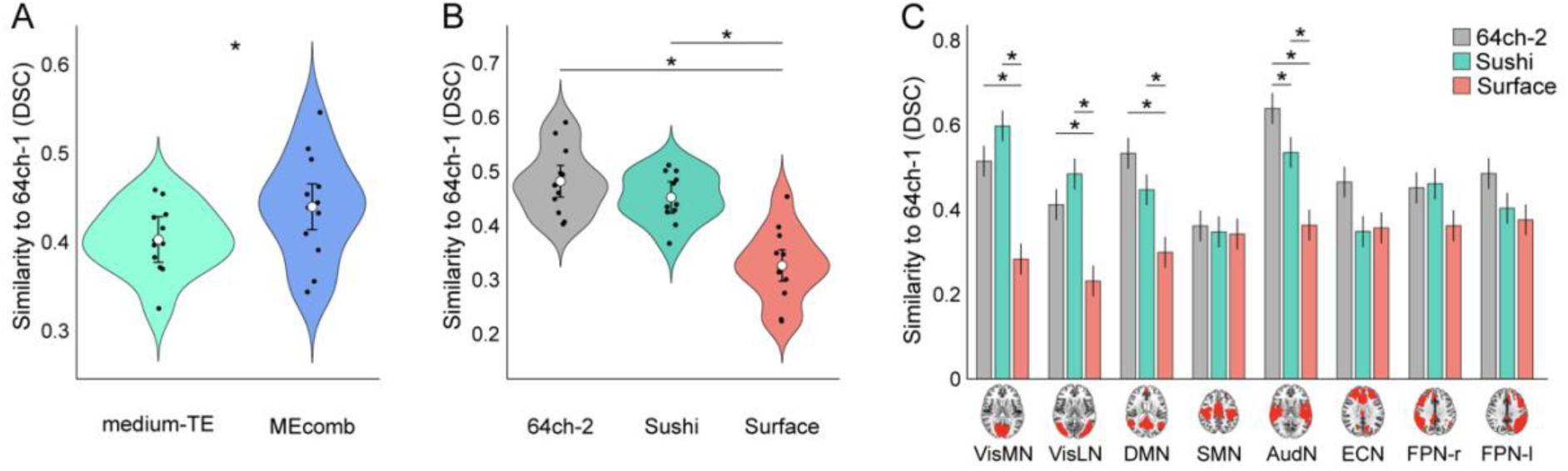
**RSN similarity**. DSC indexes the spatial similarity between subject-specific RSN maps from the 64ch-1 reference session and the comparison setups. (A) Main effect of PIPELINE, showing higher RSN similarity after MEcomb relative to the medium-TE pipeline. (B) Main effect of SETUP, showing lower RSN similarity for the Surface setup. Violins show the distribution of subject-level values, black dots show subject means, white circles show estimated marginal means, and error bars indicate 95% CIs. (C) SETUP × RSN interaction, showing that the Surface setup showed lower similarity especially for visual, default-mode, and auditory networks. Canonical RSN masks are shown below the x-axis for spatial reference. Bars show estimated marginal means, and error bars indicate standard errors. Asterisks mark significant post hoc comparisons.

Group-level RSN maps across setups are provided in Fig. S3 for qualitative visualization.

### 3.2 Experiment 2

The LMM on DSC of task-activation maps revealed a significant SETUP × PIPELINE interaction (F_2,24_ = 4.97, p = 0.016, ηp^2^ = 0.29), whereas the main effects of SETUP and PIPELINE were not significant (p ≥ 0.17). Follow-up contrasts showed that MEcomb increased DSC relative to the medium-TE pipeline for Sushi (Δ = 0.045 ± 0.015, p = 0.021), but not for 64ch-2 or Surface (p ≥ 0.45). Setup comparisons were not significant within either pipeline (p ≥ 0.38; Fig. 3).

**Figure 3.**
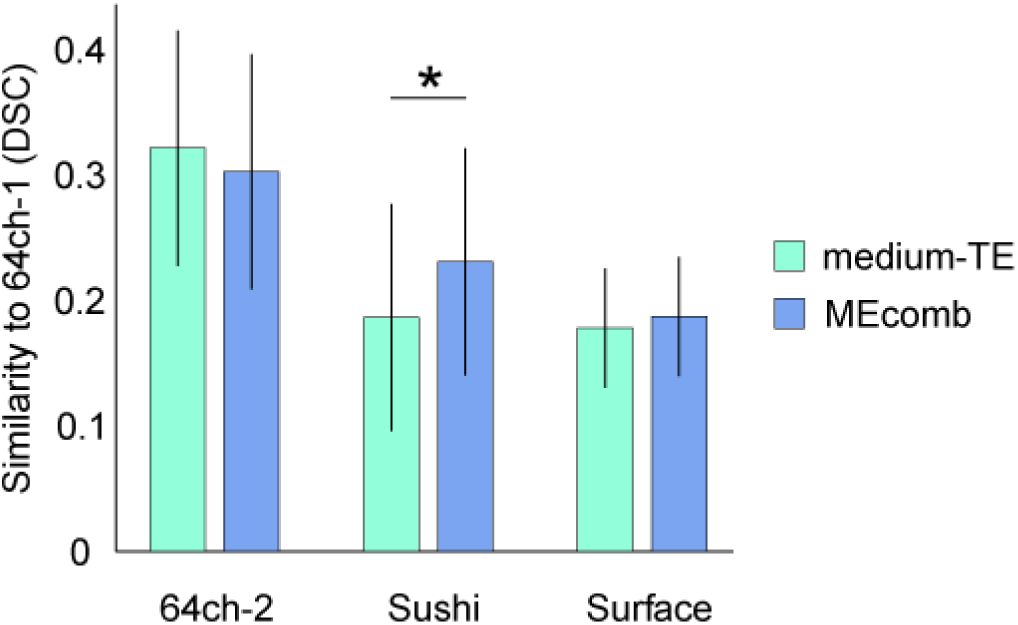
**Task-fMRI similarity**. DSC indexes the within-subject spatial similarity between task-activation maps for the 4-back > 1-back contrast obtained with the comparison setups and the corresponding 64ch-1 reference maps. Bars show estimated marginal means, and error bars indicate standard errors. MEcomb increased task-map similarity for Sushi, but not for 64ch-2 or Surface. The asterisk marks the significant post hoc comparison.

Task accuracy did not differ significantly across setups, and there was no SETUP × LOAD interaction (p ≥ 0.15), providing no evidence that the task-map similarity effects were driven by behavioral differences across setups. Accuracy was lower in the 4-back than in the 1-back condition (F_1,7_ = 6.1, p = 0.043, ηp^2^ = 0.47; Table S2).

Group-level activation maps for the 4-back > 1-back contrast in the 64ch-1 reference session showed significant clusters encompassing bilateral middle and inferior frontal gyri, the left inferior and superior parietal lobules, left precentral cortex, and left medial superior frontal, midcingulate, and dorsal anterior cingulate cortices; the medium-TE map also included the left supplementary motor area (Tables S3 and S4). This pattern captured the frontal, parietal, and medial frontal components consistently implicated in N-back working memory (Emch et al., 2019; Rottschy et al., 2012). Setup-specific group maps are shown at Z > 3 in Fig. S4 for qualitative visualization and were not used for inferential comparisons between setups or pipelines.

### 3.3 Experiment 3

The LMM on whole-brain tSNR revealed main effects of SETUP (F_3,12_ = 29.0, p < 0.001, ηp^2^ = 0.88) and PIPELINE (F_1,48_ = 699.1, p < 0.001, ηp^2^ = 0.94). The SETUP × PIPELINE interaction was also significant (F_3,48_ = 6.1, p = 0.001, ηp^2^ = 0.28). Post hoc contrasts showed that 64ch-1 and 64ch-2 did not differ significantly (p = 0.63), but both yielded substantially higher tSNR than Sushi and Surface (p < 0.001; Fig. 4A). Sushi outperformed Surface setup (p = 0.013). MEcomb significantly increased whole-brain tSNR (Δ = 18.7 ± 0.7, p < 0.001). Expressed as percentage improvement from medium-TE to MEcomb, tSNR gains were 28.8% for 64ch-1, 28.4% for 64ch-2, 49.4% for Sushi, and 39.7% for Surface (Fig. 4B).

**Figure 4.**
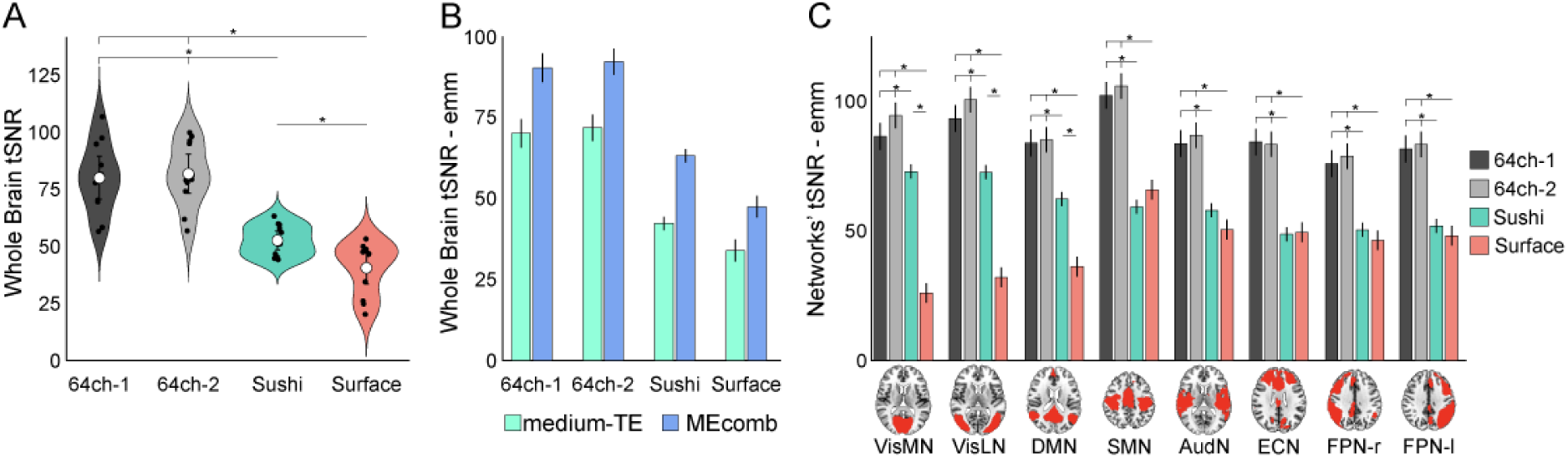
**Whole-brain and RSN-wise tSNR in the human benchmark dataset. tSNR was computed from the Experiment 1 rs-fMRI data and analyzed in Experiment 3**. (A) Whole-brain tSNR by receive setup. Violins show the distribution of subject-level values, black dots show subject means, white circles show estimated marginal means, and error bars indicate 95% CIs. (B) Whole-brain tSNR by setup and preprocessing pipeline, showing higher tSNR after MEcomb relative to the medium-TE pipeline. (C) RSN-wise tSNR by setup, averaged across preprocessing pipelines. Canonical RSN masks are shown below the x-axis for spatial reference. Bars show estimated marginal means, and error bars indicate standard errors. Asterisks mark significant post hoc comparisons.

The LMM on RSN-wise tSNR showed main effects of SETUP (F_3,12_ = 28.4, p < 0.001, ηp^2^ = 0.88), PIPELINE (F_1,720_ = 1130.2, p < 0.001, ηp^2^ = 0.61), and RSN (F_7,720_ = 81.0, p < 0.001, ηp^2^ = 0.44), as well as significant SETUP × PIPELINE (F_3,720_ = 10.1, p < 0.001, ηp^2^ = 0.04) and SETUP × RSN (F_21,720_ = 39.0, p < 0.001, ηp^2^ = 0.53) interactions. Other interactions were non-significant (p ≥ 0.057). MEcomb significantly improved RSN-wise tSNR across all setups, though the extent of benefit varied by setup, as indicated by the SETUP × PIPELINE interaction. Sushi exhibited the strongest benefit (+41.8%), followed by Surface (+33.7%), whereas gains were smaller for 64ch-1 (+21.4%) and 64ch-2 (+21.5%).

Post hoc contrasts on the main effect of setup showed that the two 64-channel coils did not differ (p = 0.43), and both outperformed Sushi and Surface setup (p ≤ 0.001). Notably, Sushi itself yielded higher tSNR than Surface setup (p ≤ 0.004). tSNR also varied across RSNs but the choice of setup played a major role; while post hoc comparisons of both reference coils did not differ in any network (p ≥ 0.09) and consistently outperformed Surface and Sushi in all RSNs (p ≤ 0.017), Sushi maintained significantly higher tSNR than Surface in visual and default mode networks (p < 0.001), not in other RSNs (p ≥ 0.22; Fig. 4C).

Average tSNR maps illustrate the spatial distribution of setup- and pipeline-related differences in the human benchmark dataset (Fig. 5).

**Figure 5.**
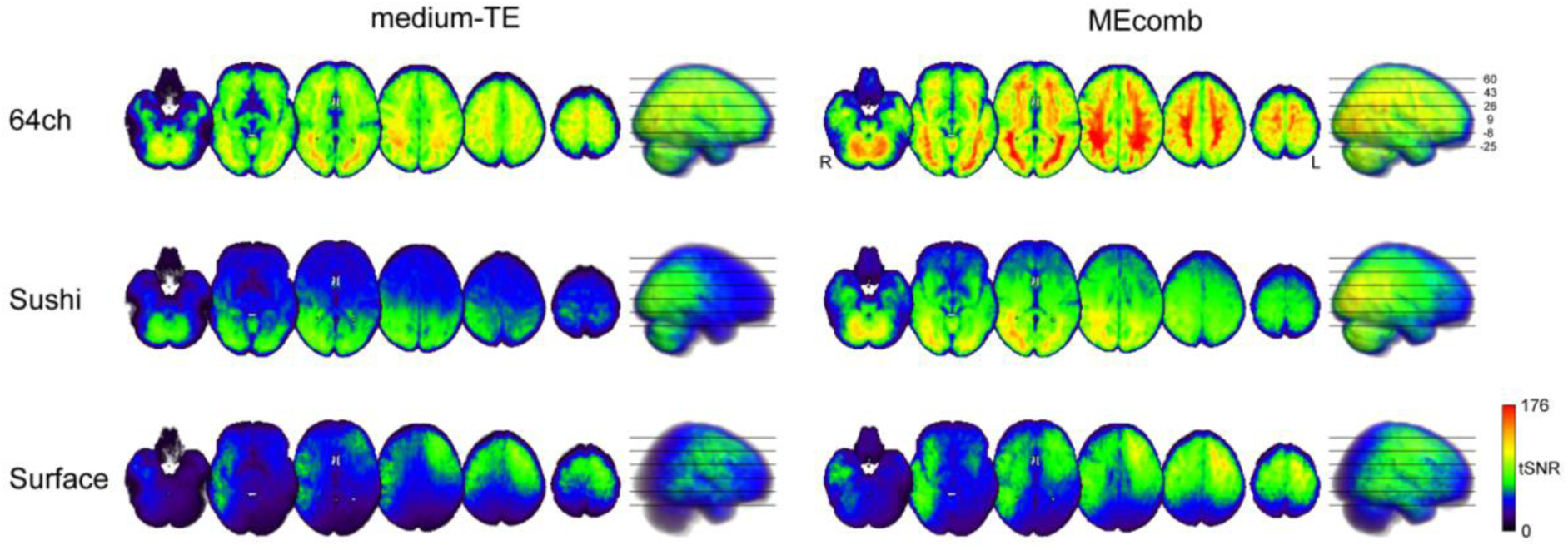
**Whole-brain tSNR in the human benchmark dataset**. Average tSNR maps from Experiment 3 are shown for each receive setup and preprocessing pipeline. Rows show the 64ch, Sushi, and Surface setups. Columns show the medium-TE and MEcomb pipelines. Axial slices and lateral renderings are displayed for spatial reference using matched slice positions within each setup. Color bars indicate tSNR.

### 3.4 Experiment 4

In the phantom benchmark sequence, MEcomb increased tSNR consistently relative to the medium-TE data. Without the TMS coil, whole-volume tSNR increased from 139.6 to 216.8, corresponding to a 55.3% gain. With the TMS coil in place, MEcomb produced comparable gains, increasing whole-volume tSNR from 129.3 to 202.5 corresponding to a 56.6% gain. Similar gains were observed in the superficial ROI (distance-to-TMS coil 25 mm; 58.3% without the TMS coil and 60.9% with the TMS coil) as well as in the deeper ROI (distance-to-TMS coil ∼75 mm; 56.4% without TMS coil and 58.4% with the TMS coil).

The passive presence of the TMS coil consistently resulted in a tSNR reduction. In the medium-TE data, whole-volume tSNR decreased from 139.6 without the TMS coil to 129.3 with the TMS coil, corresponding to a 7.4% loss. The corresponding loss was 19.5% in the superficial ROI and 6.3% in the deeper ROI. A similar pattern was observed after MEcomb (Fig. 6A). Whole-volume tSNR decreased from 216.8 to 202.5, corresponding to a 6.6% loss, while the superficial and deeper ROIs showed losses of 18.2% and 5.1%, respectively.

**Figure 6.**
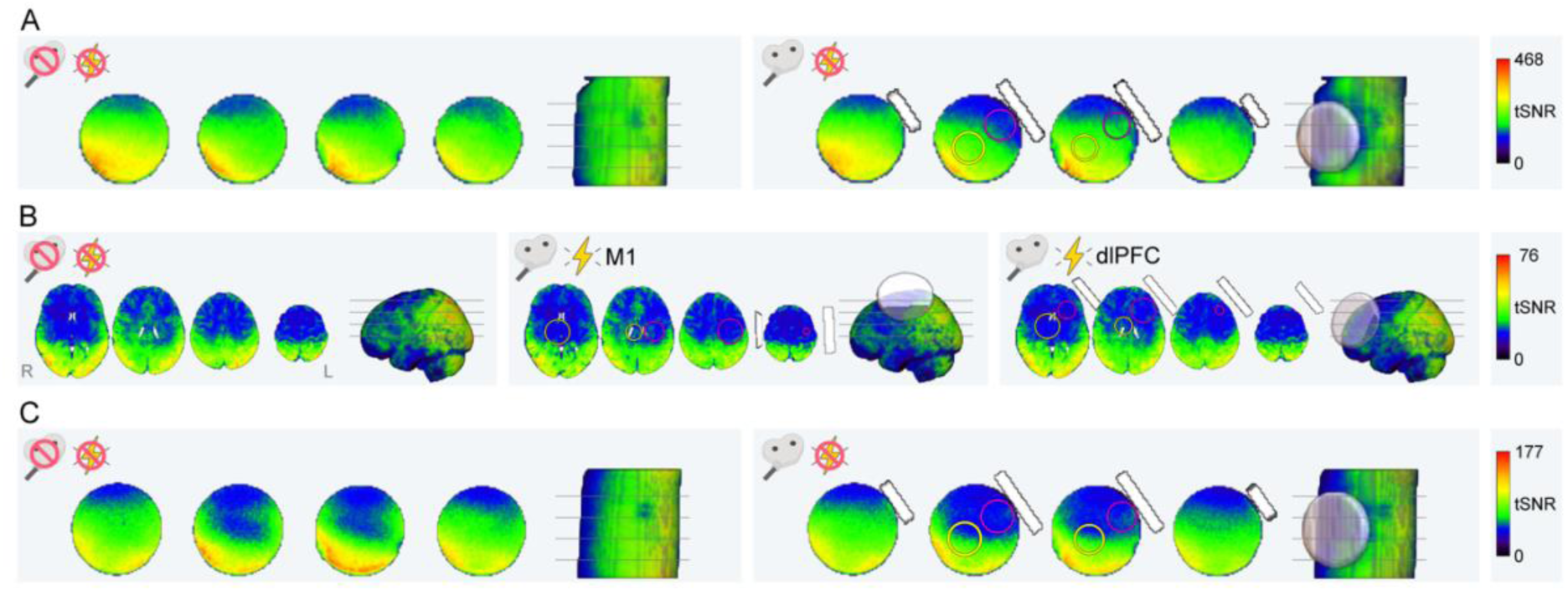
Sushi tSNR across sequences and TMS coil conditions. MEcomb tSNR maps are shown for the benchmark phantom sequence, the interleaved human sequence, and the interleaved phantom sequence. (A) Experiment 4: phantom tSNR for the benchmark sequence acquired without and with the TMS coil. (B) Experiment 5: human tSNR for the interleaved sequence acquired without TMS coil and stimulation, and during active spTMS-fMRI with the TMS coil positioned over M1 or dlPFC. (C) Experiment 6: phantom tSNR for the interleaved sequence acquired without and with the TMS coil. Axial slices and lateral renderings are shown for spatial reference. Pink and yellow circles indicate the superficial and deeper ROIs, respectively. ROIs are displayed only in the conditions with the TMS coil for visual clarity, but identical ROIs were placed in the corresponding no-coil reference data and used for the tSNR reduction comparisons. The TMS coil rendering indicates the measured coil position but is schematic and does not represent the exact shape or size of the MRI-B91 coil used in the experiments. Color scales are matched within each panel but differ across panels.

### 3.5 Experiment 5

In the active spTMS-fMRI runs, TMS pulse-related artifact screening did not reveal detectable image disturbances. For both M1 and dlPFC stimulation, no multiband slice package exceeded the predefined correlation threshold for package-level PSNR fluctuations, and no slice-level PSNR drop exceeded the predefined event threshold. Consequently, no suspicious volume, slice, or package events were identified in either run. These findings indicate that, within the sensitivity of this screening metric, delivering single TMS pulses between multiband slice packets did not produce detectable pulse-locked image artifacts in the MEcomb data.

Quantification of tSNR in the same interleaved-sequence data consistently showed that MEcomb increased absolute tSNR relative to the medium-TE data. In the no-coil/no-stimulation reference run, whole-brain tSNR increased from 15.6 to 23.9, corresponding to a 53.5% gain. During active spTMS-fMRI, whole-brain tSNR increased from 14.4 to 22.3 for M1 coil placement and from 14.9 to 23.3 for dlPFC coil placement, corresponding to gains of 54.7% and 56.6%, respectively. Thus, the tSNR benefit of MEcomb was preserved during active interleaved TMS-fMRI acquisition.

For active M1 spTMS-fMRI, whole-brain MEcomb tSNR decreased from 23.9 in the no-coil/no-stimulation reference run to 22.3, corresponding to a 6.9% reduction. The superficial M1 ROI (distance-to-TMS coil ∼50 mm) showed a 13.7% reduction while the deeper M1 ROI (distance-to-TMS coil ∼100 mm) showed a 4.2% reduction (Fig. 6B, central panel).

For active dlPFC spTMS-fMRI, whole-brain MEcomb tSNR decreased from 23.9 in the no-coil/no-stimulation reference run to 23.3, corresponding to a 2.6% reduction. The superficial dlPFC ROI (distance-to-TMS coil ∼45 mm) showed a 9.9% reduction while the deeper dlPFC ROI (distance-to- TMS coil ∼95 mm) showed a 3.8% reduction (Fig. 6B, right panel).

### 3.6 Experiment 6

The interleaved sequence tested on the phantom showed the same pattern. In the MEcomb data, whole-volume tSNR decreased from 75.6 without the TMS coil to 69.4 with the TMS coil, corresponding to an 8.2% loss. The tSNR loss was strongest in the superficial ROI (distance-to-TMS coil ∼25 mm), where values decreased by 20.2%. In the deeper ROI (distance-to-TMS coil ∼75 mm), tSNR decreased by 9.2% (Fig. 6C).

## 4 DISCUSSION

Perturbing a cortical site with TMS can alter neural activity both at the stimulation target and across connected networks (Bergmann & Hartwigsen, 2021; Ferbert et al., 1992; Momi et al., 2021). Concurrent TMS-fMRI therefore requires a BOLD readout that is not restricted to the stimulated cortex, but can sample distributed responses with sufficient sensitivity (Bergmann et al., 2021; Bergmann & Hartwigsen, 2021). The present study tested whether the TMS-compatible Sushi setup can support such a readout. The aim was not to outperform a standard 64-channel head/neck array in raw signal quality, but to provide a practical solution for concurrent TMS-fMRI: physical access for the TMS coil, whole-brain functional coverage, and temporal stability for network-level analyses.

Across six experiments, we evaluated Sushi by moving from functional readout, to signal quality, to active TMS-fMRI compatibility. Experiment 1 showed that Sushi recovered resting-state networks more similarly to the 64-channel reference than Surface, whereas Experiment 2 did not reveal significant differences between setups in task-map similarity but showed higher task-map similarity with MEcomb specifically for Sushi. Experiment 3 placed these functional findings in the context of human tSNR, where Sushi showed an intermediate but more balanced whole-brain profile than Surface. Experiments 4 and 6 quantified the tSNR cost of placing the TMS coil in the Sushi setup under controlled phantom conditions, while Experiment 5 tested the realistic concurrent TMS-fMRI with active stimulation case in one participant, showing limited target-dependent tSNR reductions and no detectable pulse-locked image artifacts. Across these metrics, MEcomb strengthened the readout relative to the conventional medium-TE pipeline. Together, these findings support Sushi and MEcomb as complementary parts of a practical solution for TMS-compatible whole-brain fMRI.

We first asked whether the whole-brain readout provided by Sushi was sufficient to identify subject-specific RSN maps. This was a critical test because canonical RSNs capture distributed functional topographies that closely correspond to the major networks recruited during task activation (Smith et al., 2009). The 64ch-1 session provided the reference pattern, and its correspondence with the canonical templates confirmed that the benchmark maps were plausible. Relative to this reference, Sushi approached the 64-channel test-retest comparison, whereas Surface showed weaker similarity, especially for networks whose spatial extent or position makes them more vulnerable to uneven receive sensitivity. This pattern is consistent with the expected receive-profile differences between the setups. Compared with Sushi, the Surface arrays likely favored regions close to the receiver elements, while more distant midline, occipital, and temporal nodes were sampled with lower and less homogeneous sensitivity. This may explain why Sushi recovered the default-mode, visual, and auditory networks more faithfully than Surface in the present array configuration.

By reliably engaging a distributed working-memory network, the N-back task provided a complementary benchmark of task-evoked BOLD responses (Emch et al., 2019; Rottschy et al., 2012). The 4-back > 1-back maps obtained with the 64ch-1 reference showed the expected frontal, parietal, and medial frontal engagement, consistent with previous letter N-back fMRI studies (Clark et al., 2017; Miri Ashtiani & Daliri, 2023). Task-map similarity did not differ significantly between setups within either pipeline and therefore did not establish superiority of one receive setup. Instead, the interaction reflected higher DSC for Sushi when the Sushi and 64ch-1 maps were both processed with MEcomb than when both were processed as medium-TE images. DSC did not differ significantly between pipelines for either the 64ch-2-64ch-1 or Surface-64ch-1 comparison. Because DSC was calculated from thresholded single-subject maps, it provided a stringent test of within-subject spatial correspondence. Robust group-level activation does not necessarily imply equally stable individual-level maps, particularly given the limited test-retest reliability often observed for individual-level task-fMRI measures (Elliott et al., 2020). Moreover, because DSC penalizes both missing reference activation and additional non-overlapping activation, similar DSC values may reflect different spatial profiles. Consistent with this, the qualitative group maps appeared more spatially restricted for Sushi and more widespread for Surface, although these maps were not used for inferential comparisons. This type of functional benchmark is directly relevant to online TMS-fMRI, where task-evoked BOLD activity is measured while TMS manipulates a network node, allowing its effects across the engaged network to be assessed (Grosshagauer et al., 2024; Schuler & Hartwigsen, 2025).

The predominantly lateral frontal and parietal distribution of the N-back network, overlapped well with the regions physically covered by the Surface arrays. The task may therefore have been a favorable benchmark for targeted Surface reception. The RSNs, by contrast, included more widely distributed posterior, temporal, and midline components that were particularly vulnerable to uneven receive sensitivity, which may explain the clearer separation between Sushi and Surface. Experiment 3, accordingly, showed high local tSNR near the Surface arrays, but also demonstrated that this sensitivity depended strongly on their placement. Sushi did not provide the same local receiver advantage, but offered a more balanced whole-brain profile. Despite not reaching the absolute tSNR of the 64-channel reference, as expected for a TMS-compatible setup, Sushi provided higher whole-brain and RSN-wise tSNR than Surface. Across all three setups, MEcomb further increased whole-brain tSNR relative to the medium-TE image, with gains of ∼40-50% for the TMS-compatible setups and ∼28% for the 64-channel array.

Crucially, Experiment 5 provided a proof-of-principle test of Sushi compatibility with active interleaved spTMS-fMRI. The interleaved BOLD-EPI sequence was designed to deliver single TMS pulses between multiband slice packets, leaving approximately 30 ms between pulse delivery and the onset of the next packet. Under these timing conditions, artifact screening did not reveal detectable pulse-locked image disturbances, and tSNR measurements showed that the interleaved acquisition remained usable during active stimulation. These findings indicate that, under the tested conditions, Sushi can support active interleaved spTMS-fMRI without evident pulse-locked image artifacts.

The comparisons conducted in the present study help define the use cases in which each TMS-compatible setup may be preferable. Sushi is particularly suited to questions requiring spatially distributed or not fully predictable network-level readouts, such as mapping remote effects beyond the stimulated cortex. In these cases, its more balanced whole-brain profile is advantageous, even though it does not provide the same local receiver advantage as a surface array placed directly over the target. Surface remains valuable when the primary aim is to maximize sensitivity near the TMS coil or across a limited set of predefined regions. It also leaves more free space around the head in the bore than Sushi, which can improve the line-of-sight for neuronavigation and facilitate TMS-coil placement during preparation. Both TMS-compatible setups were rated as generally comfortable by participants, although Surface tended to have lower comfort ratings (see Supplementary Materials).

The same flexibility that makes Surface useful for targeted applications can, however, complicate network-level comparisons. In our experiment, the TMS coil and the first surface array were mounted together and centered over the intended target region, either left motor or left prefrontal cortex, while the second surface array was positioned to extend coverage. In routine concurrent TMS-fMRI, this second array is often positioned according to the hypothesized downstream regions of interest, for example over contralateral, prefrontal, or occipital areas. This targeted placement can be advantageous, but it also means that the regions sampled with high sensitivity depend on array placement and may differ across targets, participants, and experimental conditions. This is relevant for distributed systems such as working-memory networks, whose recruited nodes may vary across tasks, individuals, or groups (Emch et al., 2019; Yaple et al., 2019). This issue becomes particularly critical in studies comparing different stimulation sites, because shifting the TMS target requires repositioning the primary Surface array, which is mounted on the TMS coil, and may also require repositioning the second array. As a result, changes in the coverage pattern across conditions may partly affect the apparent functional readout, rather than reflecting TMS-induced neural effects alone. In the present study, motor and prefrontal placements were pooled to evaluate functional readouts that were not confined to the immediate vicinity of the stimulation target. This choice may have increased variability for the Surface setup, but it also provided a realistic test of its suitability for network-level applications. Overall, the RSN and tSNR findings favor Sushi when stable, broadly distributed whole-brain coverage is required, whereas the task benchmark suggests that Surface can provide an adequate functional readout when its sensitivity profile overlaps the principal network nodes, although it did not outperform Sushi.

The different strengths of Sushi and Surface are also linked to how the receive elements are positioned relative to the scalp and the TMS coil. In the Surface setup, the local receive array lies between the scalp and the TMS coil. This geometry is advantageous for local imaging, as shown by the strong local SNR and tSNR reported for TMS-dedicated surface arrays. Navarro de Lara et al. (2015) reported a 5-fold SNR gain at approximately 3 cm depth relative to a birdcage setup, based on 3D GRE SNR maps. Jackson et al. (2024) similarly showed very high EPI tSNR directly beneath TMS-dedicated surface arrays. However, this arrangement increases the effective coil-to-scalp distance by 4.5 mm, thereby reducing stimulation efficiency. Navarro de Lara et al. (2015) measured a TMS-efficiency loss of 14-16% in phantom and observed an increase in motor threshold from 66% to 84% MSO in one participant when the array was placed between the TMS coil and the head. A similar distance-related cost was reported for the RF Cap, where the additional 3-4 mm thickness increased resting motor threshold (Navarro De Lara et al., 2025). This trade-off can matter in participants with high thresholds and in longer protocols, where higher stimulator output may accelerate coil heating. Sushi avoids placing a local receive array between the TMS coil and scalp, but at the cost of lower local receive sensitivity than a surface array positioned directly over the target.

Previous TMS-compatible hardware studies have shown that the TMS coil can reduce SNR or tSNR nearby or alter the spatial sensitivity profile when it is physically present during imaging (Jackson et al., 2024; Navarro De Lara et al., 2025). For example, Navarro De Lara et al. (2025) reported a 5-20% local SNR loss when the TMS coil was placed over the RF Cap, based on GRE SNR maps in a phantom. Direct comparison with our data requires caution because GRE SNR maps provide a controlled estimate of static image SNR and receive-sensitivity changes, whereas EPI tSNR captures the temporal stability of the functional time series and can also reflect EPI-specific effects such as susceptibility-related signal loss, dropout, geometric distortion, reconstruction choices, and scanner-related temporal instability (Jamil et al., 2021; Olman et al., 2009; Welvaert & Rosseel, 2013). We quantified this practical TMS coil cost in Sushi using the same multi-echo BOLD-EPI sequences used in the study. In Experiment 4, using the benchmark sequence, placing the TMS coil against the phantom reduced MEcomb tSNR by 6.6% at the whole-volume level and by 18.2% in the superficial ROI close to the TMS coil. In Experiment 6, using the interleaved sequence optimized for active interleaved TMS-fMRI, MEcomb tSNR decreased by 8.2% at the whole-volume level, and by 20.2% in the superficial ROI. Experiment 5 then measured the same interleaved sequence in a human participant during active concurrent spTMS-fMRI. Relative to the no-coil/no-stimulation reference, whole-brain MEcomb tSNR reductions were limited for both M1 (6.9%) and dlPFC (2.6%) coil placements. Local reductions were target-dependent, with the superficial ROI reduced by 13.7% for the M1 target and 9.9% for the dlPFC target. These results show that the TMS coil introduced a measurable but spatially graded tSNR cost in Sushi. This pattern is consistent with previous TMS-MRI work showing local signal loss and geometric distortion in T2*-weighted EPI near MR-compatible TMS coils, with weaker effects in humans because the cortex is separated from the coil by scalp, skull, and other non-brain tissue (Baudewig et al., 2000; Mizutani-Tiebel et al., 2022; Riddle et al., 2022). Accordingly, losses were smaller in the superficial ROI for the human than the phantom, consistent with the greater distance between the TMS coil and the nearest cortex. The observed losses are also compatible with TMS-compatible RF-array work showing that TMS coil placement can produce local SNR reductions and interact with nearby receive elements (Navarro de Lara et al., 2015; Navarro De Lara et al., 2025). In addition to this passive coil-placement cost, active TMS-fMRI can introduce further artifact sources, including RF noise carried through the TMS system and residual leakage currents through the stimulation coil, which can reduce EPI signal stability or produce local EPI signal changes if not properly controlled (Bungert et al., 2012; Weiskopf et al., 2009). Because we did not separately measure B0, B1+, RF noise, or receive-element detuning, these mechanisms should be interpreted as plausible contributors rather than isolated causes of the observed tSNR loss. Thus, the phantom measurements provide a conservative estimate close to the TMS coil, while the human data indicate that the practical tSNR cost in cortical tissue was smaller under realistic active TMS-fMRI conditions.

MEcomb consistently increased tSNR relative to the conventional medium-TE image across setups and sequences. Multi-echo EPI samples the BOLD signal at multiple TEs and allows the echo time series to be combined according to their TE-dependent signal properties (Kundu et al., 2017; Posse et al., 1999). This does not mean that MEcomb increases raw SNR relative to the shortest TE image, which usually has the highest signal amplitude. Rather, in this study MEcomb improved temporal stability relative to the medium-TE image commonly used for BOLD-sensitive single-echo fMRI. This benefit was observed in the 64-channel, Sushi, and Surface setups, in both human and phantom data, and in the interleaved sequence used for active spTMS-fMRI. The practical trade-off is that multi-echo acquisition requires additional echo sampling, which can increase TR unless other parameters such as spatial resolution, coverage, or acceleration are adjusted. In 3T protocols, the TR cost is quantifiable on the order of 20-30%, depending on number of echoes, their timing, and acceleration (Kovářová et al., 2022). In this study, the minimum TR increased by 27% when moving from a single-echo acquisition at the medium TE to the three-echo benchmark protocol. MEcomb therefore acted as a processing complement to the Sushi hardware: Sushi addressed the receive-geometry constraint imposed by TMS access, while MEcomb improved the temporal stability and comparability of the resulting fMRI time series. Its benefit was not specific to Sushi, but its practical relevance is greatest for TMS-compatible acquisitions, where receive geometry is necessarily constrained by the need to position the TMS coil.

In Experiments 1 and 2 we did not deliver TMS pulses during fMRI; however, TMS coil placement, filtering, and hardware were configured for concurrent operation, but the present datasets evaluate functional validity without on-line perturbation. This design focused on how receive geometry and post-hoc optimization jointly affect the fidelity of the fMRI readout. Active TMS-fMRI compatibility was tested in a single participant and with single-pulse stimulation only in Experiment 5. This experiment was designed as a technical feasibility and artifact-control test, not as a full physiological TMS-fMRI study. We therefore did not model BOLD responses to stimulation. Nevertheless, the absence of detectable pulse-locked artifacts, together with limited tSNR reductions during active acquisition, provides proof-of-principle evidence that the Sushi setup can support active interleaved spTMS-fMRI under the tested conditions. Future studies should extend this validation to larger samples, additional targets, repeated-session reliability, and patterned stimulation protocols.

The Sushi setup should be interpreted in the context of ongoing efforts to improve TMS-compatible MR receive hardware. Dedicated surface-array solutions increased local sensitivity and enabled accelerated imaging compared with earlier birdcage approaches (Navarro de Lara et al., 2015), but their spatially uneven receive profile can limit studies targeting distributed network responses (Jackson et al., 2024). Newer flexible multi-channel concepts, including the RF Cap and Taco setups, further show that TMS-fMRI hardware is moving toward higher-channel, more flexible, and more whole-brain-oriented solutions (Assem et al., 2025; Navarro De Lara et al., 2025; Tik et al., 2025). Sushi follows the same general direction, but with a different practical emphasis. It repurposes readily commercially available body-array elements into a low-cost configuration that can be implemented without building a dedicated RF coil. Its main value is therefore not technical novelty at the element-design level, but a usable compromise between whole-brain coverage, TMS access, and deployability.

Taken together, the functional, human tSNR, phantom, and active interleaved acquisitions point to the same conclusion. Sushi does not reproduce the raw signal performance of a close-fitting 64-channel head/neck array, but it provides a more balanced whole-brain readout than Surface while preserving physical access for the TMS coil. Across these tests, MEcomb strengthened the readout: it increased tSNR relative to the medium-TE image, improved RSN similarity, increased task-map correspondence for Sushi, and remained beneficial in the interleaved acquisition used for active spTMS-fMRI. We therefore view Sushi and MEcomb as complementary parts of the same practical solution. Sushi addresses the hardware constraint of concurrent TMS-fMRI by providing a balanced whole-brain readout, whereas MEcomb improves the temporal stability of the acquired data. This combination is especially relevant for network-level TMS-fMRI, where the scientific target is often not only the stimulated cortex, but also distributed connected regions.

## DATA AND CODE AVAILABILITY

Analysis code is available at https://github.com/echiap/2026_Sushi_TMS-fMRI_benchmark. Due to ethics and data-protection constraints, the MRI data are not publicly available. Further information can be requested from the corresponding author.

## AUTHOR CONTRIBUTIONS

Y.X.: Investigation, Software, Formal analysis, Data Curation, Visualization, Writing – Original Draft, Writing – Review & Editing; M.B.: Methodology, Software, Resources, Writing – Review & Editing; L.M.: Investigation, Writing – Review & Editing; K.T.: Investigation, Writing – Review & Editing; M.L.: Methodology, Writing – Review & Editing; T.O.B.: Methodology, Writing – Review & Editing; M.A. N.: Conceptualization, Resources, Writing – Review & Editing; E.G.: Conceptualization, Methodology, Supervision, Resources, Project administration, Writing – Review & Editing; E.C.: Methodology, Software, Formal analysis, Visualization, Supervision, Project administration, Writing – Original Draft, Writing – Review & Editing.

## DECLARATION OF AI-ASSISTED TECHNOLOGIES

During the preparation of this manuscript, the authors used ChatGPT (GPT-5 family; OpenAI) to support language editing, manuscript organization, code drafting, and code debugging. All scientific content, analyses, code, outputs, references, and final text were critically reviewed, verified, and approved by the authors, who take full responsibility for the content of the manuscript.

## ACKNOWLEDGEMENTS

We thank Dr. Mosayebi Samani for his contributions during the early development of the Sushi coil concept. We also thank Mr. Martin Schmitz and Mr. Timo Scholz from IfADo workshop for designing, manufacturing, and iteratively refining the Sushi coil holder prototype.

## SUPPLEMENTARY MATERIAL

Supplementary material for this article is available online.

