## Supplementary Materials for "Optimizing Network-Level TMS-fMRI: Benchmarking the TMS-Compatible Sushi MR Setup"

#### 1 Supplementary methods and quality control

##### 1.1 Intensity homogenization and registration quality control

For the human benchmark datasets (Experiments 1-3), intensity homogenization was applied to reduce spatial receive-sensitivity bias before anatomical registration and downstream analysis. The correction was computed from two 2D-GRE reference images acquired with the integrated scanner body coil and with the corresponding receive setup. Both reference images were Gaussian-smoothed ($\sigma$ = 5 voxels), and a smooth correction field was obtained from the ratio between the body-coil and receive-coil images. This correction field was aligned to the EPI data and applied to each EPI volume.

This step was particularly relevant for the Surface setup, where the receive arrays were positioned laterally and produced strong spatial intensity gradients in the uncorrected EPI images. In some datasets, these gradients impaired intensity-based EPI-to-T1 registration, leading to anatomically implausible alignments. After homogenization, the EPI intensity profile was more anatomically plausible and EPI-to-T1 registration was more robust (Fig. S1).

Because the correction field was constant over time, homogenization rescaled voxel-wise signal amplitudes without altering the temporal structure of the time series. The procedure was therefore used to improve spatial intensity profiles and anatomical alignment.

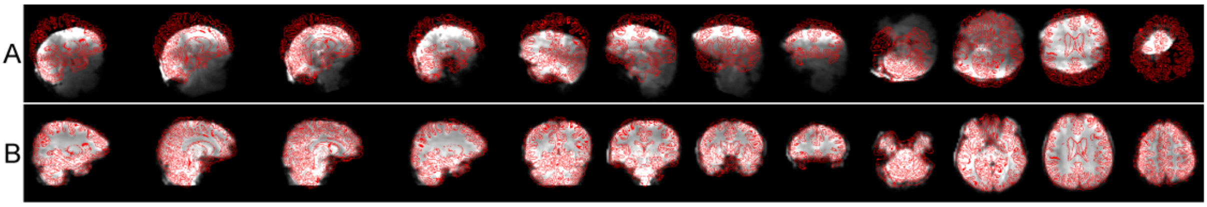


**Figure S1. Example EPI-to-T1 registration before and after homogenization.** (A) Raw Surface setup EPI registered to the participant's T1-weighted image. (B) The same registration using the homogenized EPI. Red contours show the T1 outline overlaid on the EPI. Registration was performed with rigid-body alignment (6 DOF).

##### 1.2 Functional processing details

For rs-fMRI, single-subject ICA was performed separately for each participant, setup, and preprocessing pipeline (medium-TE, MEcomb) using FSL MELODIC. IC spatial maps were registered to MNI space and compared with the selected Smith et al. (2009) RSN templates. The component-selection procedure was designed to identify subject-specific maps corresponding to the eight canonical RSNs used in the main analysis. ICs with low spatial correspondence to all selected templates were discarded. The remaining ICs were visually inspected for obvious non-canonical or artifactual spatial patterns. Components were removed only when all three independent inspectors (E.C., Y.X., M.B.) classified them as noise. The retained IC maps were then entered into the template-based selection procedure described in the main text.

For task-fMRI, each participant, setup, and preprocessing pipeline was analyzed with a first-level block-design GLM in SPM12. Each run was modeled as a separate session. Regressors for 1-back, 3-back, and 4-back blocks were convolved with the canonical HRF, and a 108 s high-pass filter was applied. The contrast of interest was 4-back > 1-back. The resulting first-level t-statistic maps were converted to Z maps, thresholded at $Z > 3$, binarized, and used for DSC analyses against the corresponding 64ch-1 reference maps within participant. At the group level, first-level contrast images were entered into a flexible factorial model including SUBJECT, SETUP (64ch-1, 64ch-2, Sushi, Surface), and PIPELINE (medium-TE, MEcomb). Cell-specific contrasts were used to estimate group activation separately for each setup and pipeline. Inferential cluster results are reported only for the 64ch-1 reference maps (Tables S3 and S4), whereas all setup-specific maps are shown for qualitative visualization (Fig. S4).

### 2 Resting-state benchmark

#### 2.1 Validation of 64ch-1 RSN reference

To quantify the correspondence between the 64ch-1 reference session and canonical RSNs, we computed subject-specific spatial correlations between RSN maps from the 64ch-1 session and the RSN templates from Smith et al. (2009). For each subject, RSN, and preprocessing pipeline (medium-TE, MEcomb), FSL fslcc was used to calculate Pearson's r between the subject-level ICA map and the corresponding RSN template within a whole-brain mask. Table S1 reports the mean r and standard deviation across subjects. Correlations ranged from 0.35 to 0.58 across individual networks. The average correlation across RSNs was 0.43 ± 0.08 for the medium-TE pipeline and 0.44 ± 0.10 for MEcomb, indicating stable and plausible correspondence between the 64ch-1 maps and the canonical RSNs.

To illustrate this correspondence, we computed a group ICA with FSL MELODIC on the 64ch-1 MEcomb data and overlaid the resulting spatial maps with the RSN templates from Smith et al. (2009). As shown in Fig. S2, the 64ch-1 networks recovered the expected topography of the selected RSNs. Together with the spatial-correlation analysis, this supports the use of 64ch-1 as the functional benchmark for comparing RSN-map similarity across setups.

Table S1. 64ch-1 and canonical RSN correlation

| | VisMN | VisLN | DMN | SMN | AudN | ECN | FPN-r | FPN-l | Mean |
|---|---|---|---|---|---|---|---|---|---|
| 64ch-1 medium-TE | 0.57 ± 0.08 | 0.46 ± 0.08 | 0.49 ± 0.08 | 0.35 ± 0.08 | 0.44 ± 0.08 | 0.36 ± 0.08 | 0.42 ± 0.08 | 0.40 ± 0.08 | 0.43 ± 0.08 |
| 64ch-1 MEcomb | 0.58 ± 0.10 | 0.48 ± 0.10 | 0.50 ± 0.10 | 0.36 ± 0.10 | 0.46 ± 0.10 | 0.37 ± 0.10 | 0.41 ± 0.10 | 0.41 ± 0.10 | 0.44 ± 0.10 |

Spatial correlation (Pearson's r) between 64ch-1 RSN maps and Smith et al. 2009 templates, by pipeline. Values are mean ± SD across subjects.

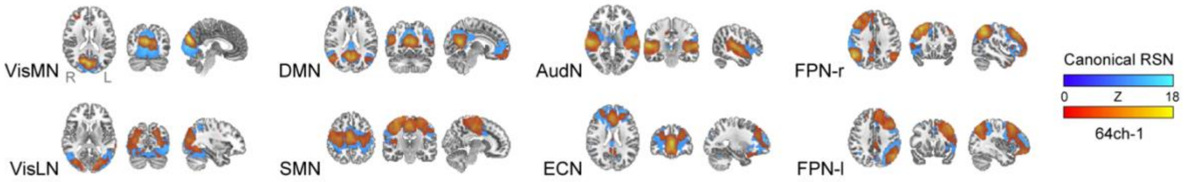


**Figure S2. Group-ICA RSN maps for the 64ch-1 reference session.** Group-level rs-fMRI ICA maps are shown for illustrative purposes only. The panels illustrate, for each RSN, the qualitative overlap between the 64ch-1 MEcomb maps and the canonical RSN templates from Smith et al. (2009).

#### 2.2 Group-ICA RSN maps

Group-ICA maps were computed with FSL MELODIC separately for each setup and preprocessing pipeline to illustrate typical network topographies (Fig. S3). These group-level maps were used for visualization only.

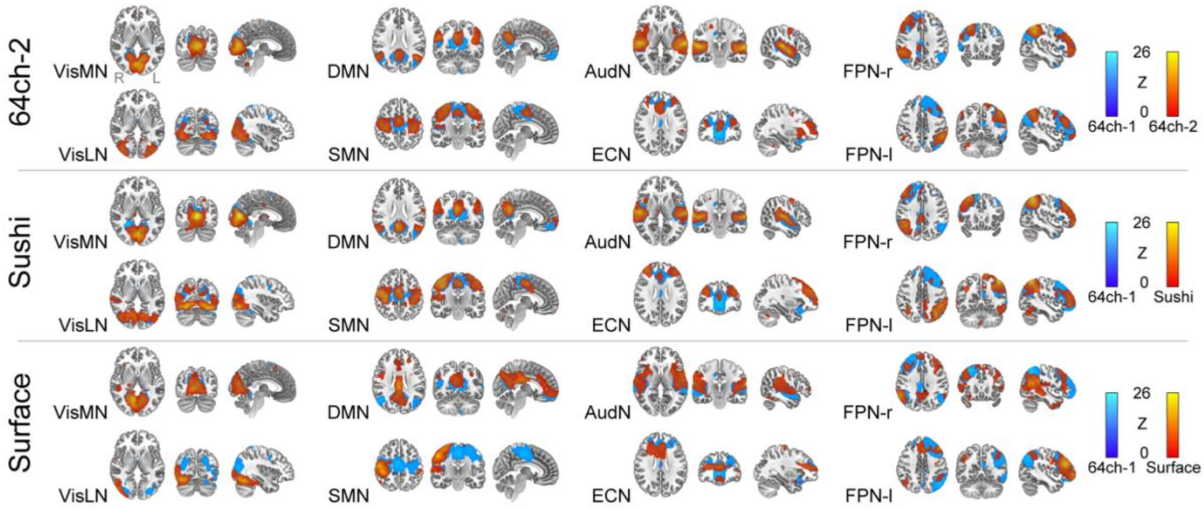


**Figure S3. Group-ICA RSN maps for the tested setups.** Group-level maps are thresholded at Z > 3. Cool colors show the 64ch-1 reference maps, and overlapping warm colors show the corresponding maps from each comparison setup.

### 3 Task-fMRI benchmark

#### 3.1 N-back task performance

To evaluate whether behavioral performance could account for differences in 4-back > 1-back activation-map similarity, accuracy was compared across setups. In the task-fMRI subset (n = 8), accuracy was analyzed with a repeated-measures ANOVA with within-subject factors SETUP (64ch-1, 64ch-2, Sushi, Surface) and LOAD, where LOAD refers to N-back difficulty (1-back, 4-back). The main effect of LOAD ($F_{1,7} = 6.1$, $p = 0.043$, $\eta p^2 = 0.47$) indicates lower accuracy in the 4-back than in the 1-back condition (Table S2). Accuracy did not differ significantly across setups (Greenhouse-Geisser corrected: p = 0.15), and there was no SETUP × LOAD interaction (p = 0.31).

**Table S2. N-back task performance**

| | 64ch-1 | 64ch-2 | Sushi | Surface |
|---|---|---|---|---|
| 1-back | 93.3 ± 5.7 | 89.6 ± 9.2 | 91.8 ± 5.8 | 93.6 ± 4.3 |
| 4-back | 77.5 ± 21.9 | 68.3 ± 24.3 | 71.9 ± 23.6 | 77.7 ± 23.2 |

Values represent mean ± SD accuracy in percent across participants for each setup and LOAD level. Accuracy was calculated across target and non-target trials, with omissions treated as incorrect responses.

#### 3.2 Validation of the 64ch-1 N-back task activation pattern

For the 64ch-1 reference session, we report group-level clusters for the 4-back > 1-back contrast that survived cluster-level FWE correction (cluster-forming threshold: p < 0.001 uncorrected; cluster-level threshold: pFWE < 0.05). This analysis was performed to document that the reference setup recovered the expected working-memory activation pattern. Results are reported separately for the medium-TE pipeline (Table S3) and MEcomb pipeline (Table S4). Across pipelines, significant clusters encompassed bilateral middle and inferior frontal gyri, the left inferior and superior parietal lobules, left precentral cortex, and left medial superior frontal, midcingulate, and dorsal anterior cingulate cortices; the medium-TE map also included the left supplementary motor area. These group-level maps were used only to validate the reference activation pattern and for qualitative visualization. No inferential comparisons between setups or preprocessing pipelines were performed on these group-level cluster results.

**Table S3. fMRI results of task analysis with 64ch-1 medium-TE**

| Cluster ID | Cluster pFWE | Cluster Size | Cluster Label | Cluster Label % | Peak Z | Peak MNI Coordinates x | y | z | Peak Label |
|---|---|---|---|---|---|---|---|---|---|
| 1 | < 0.001 | 5417 | 'Frontal_Inf_Tri_L' | 47.3 | 6.22 | -44 | 26 | 14 | 'Frontal_Inf_Tri_L' |
| | | | 'Frontal_Mid_2_L' | 20.3 | 5.65 | -50 | 21 | 16 | 'Frontal_Inf_Tri_L' |
| | | | 'Frontal_Inf_Oper_L' | 15.4 | 5.36 | -40 | 3 | 33 | 'Precentral_L' |
| | | | 'Precentral_L' | 12.2 | | | | | |
| 2 | < 0.001 | 3232 | 'Parietal_Inf_L' | 63.8 | 5.57 | -46 | -40 | 40 | 'Parietal_Inf_L' |
| | | | 'Parietal_Sup_L' | 27.2 | 5.32 | -32 | -57 | 45 | 'Parietal_Inf_L' |
| | | | 'OUTSIDE' | 7.8 | 4.98 | -44 | -48 | 48 | 'Parietal_Inf_L' |

| | | | | | | | | | |
|---|---|---|---|---|---|---|---|---|---|
| 3 | 0.01 | 511 | 'Frontal_Sup_Medial_L' | 63.6 | 4.74 | -6 | 28 | 33 | 'Frontal_Sup_Medial_L' |
| | | | 'Cingulate_Mid_L' | 20.7 | 3.61 | -3 | 21 | 45 | 'Supp_Motor_Area_L' |
| | | | 'Supp_Motor_Area_L' | 7.4 | | | | | |
| | | | 'ACC_sup_L' | 6.8 | | | | | |
| 4 | 0.004 | 618 | 'Frontal_Mid_2_R' | 71.2 | 4.41 | 34 | 10 | 62 | 'Frontal_Mid_2_R' |
| | | | 'Frontal_Sup_2_R' | 10.2 | 3.85 | 45 | 16 | 51 | 'Frontal_Mid_2_R' |
| | | | 'OUTSIDE' | 10.0 | 3.82 | 48 | 15 | 39 | 'Frontal_Mid_2_R' |
| | | | 'Frontal_Inf_Oper_R' | 8.3 | | | | | |

Clusters showing greater activation for the 4-back > 1-back contrast in the 64ch-1 medium-TE data. Effects were tested using a voxelwise cluster-forming threshold of $p < 0.001$ (uncorrected) and cluster-level pFWE < 0.05. AAL3 regions comprising at least 5% of each cluster are reported, together with cluster size (k), cluster-level pFWE, peak Z values, and peak MNI coordinates (x, y, z).

Table S4. fMRI results of task analysis with 64ch-1 MEcomb

| Cluster ID | Cluster pFWE | Cluster Size | Cluster Label | Cluster Label % | Peak Z | Peak MNI Coordinates | | | Peak Label |
|---|---|---|---|---|---|---|---|---|---|
| | | | | | | x | y | z | |
| 1 | < 0.001 | 4768 | 'Frontal_Inf_Tri_L' | 50.1 | 5.78 | -45 | 24 | 14 | 'Frontal_Inf_Tri_L' |
| | | | 'Frontal_Mid_2_L' | 19.8 | 5.22 | -40 | 4 | 33 | 'Precentral_L' |
| | | | 'Frontal_Inf_Oper_L' | 15.3 | 5.18 | -51 | 14 | 20 | 'Frontal_Inf_Oper_L' |
| | | | 'Precentral_L' | 11.4 | | | | | |
| 2 | < 0.001 | 2832 | 'Parietal_Inf_L' | 63.7 | 5.13 | -46 | -40 | 40 | 'Parietal_Inf_L' |
| | | | 'Parietal_Sup_L' | 26.3 | 5.13 | -32 | -57 | 45 | 'Parietal_Inf_L' |
| | | | 'OUTSIDE' | 8.2 | 4.87 | -44 | -46 | 46 | 'Parietal_Inf_L' |
| 3 | 0.033 | 395 | 'Frontal_Sup_Medial_L' | 62.3 | 4.38 | -6 | 27 | 33 | 'Cingulate_Mid_L' |
| | | | 'Cingulate_Mid_L' | 28.4 | | | | | |
| | | | 'ACC_sup_L' | 7.8 | | | | | |
| 4 | 0.003 | 657 | 'Frontal_Mid_2_R' | 67.6 | 4.32 | 36 | 12 | 60 | 'Frontal_Mid_2_R' |
| | | | 'Frontal_Inf_Oper_R' | 14.8 | 4.16 | 52 | 16 | 42 | 'Frontal_Mid_2_R' |
| | | | 'OUTSIDE' | 11.1 | 4.02 | 45 | 16 | 52 | 'Frontal_Mid_2_R' |

Clusters showing greater activation for the 4-back > 1-back contrast in the 64ch-1 MEcomb data. Effects were tested using a voxelwise cluster-forming threshold of $p < 0.001$ (uncorrected) and cluster-level pFWE < 0.05. AAL3 regions comprising at least 5% of each cluster are reported, together with cluster size (k), cluster-level pFWE, peak Z values, and peak MNI coordinates (x, y, z).

#### 3.3 Group-level N-back task-activation maps

Setup-specific group-level activation maps for the 4-back > 1-back contrast are shown in Fig. S4 for qualitative visualization.

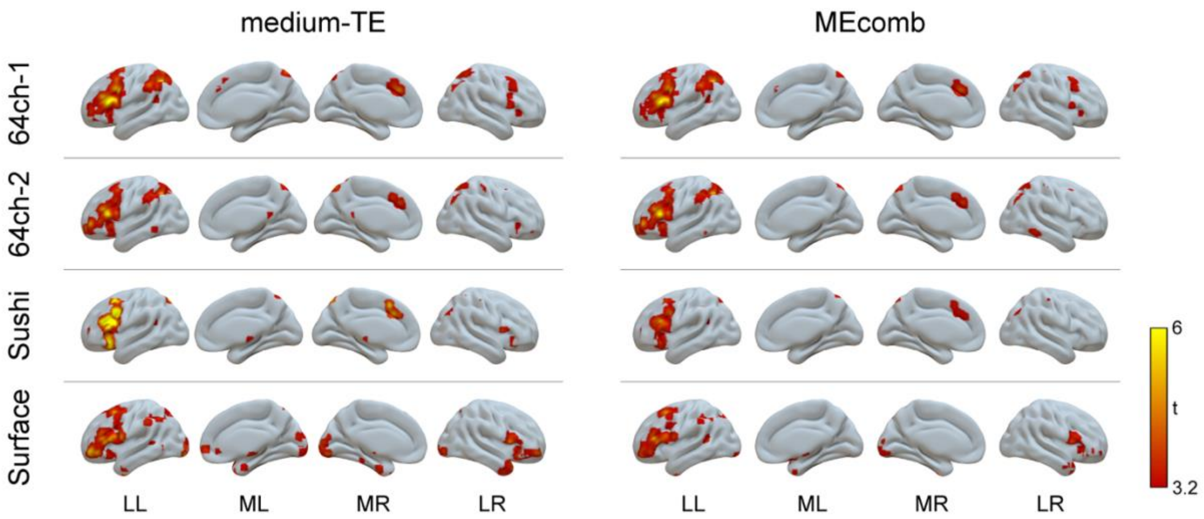

**Figure S4. Group-level task-activation maps for Experiment 2.** Maps show the 4-back > 1-back contrast for each setup and preprocessing pipeline. Maps are displayed at T > 3.16 (corresponding to Z > 3), without an extent threshold, using a common display range across panels. They are shown for qualitative visualization only and were not entered into inferential setup or pipeline comparisons. LL = lateral view, left hemisphere; ML = medial view, left hemisphere; MR = medial view, right hemisphere; LR = lateral view, right hemisphere.

#### 4 Subjective comfort ratings

Participants rated session comfort on a 5-point Likert scale after each scan (1 = very comfortable, 5 = not comfortable at all). For the 64-channel receive coil, participants provided a single comfort rating covering both 64ch sessions. Comfort ratings were therefore compared across three setup levels (64ch, Sushi, Surface; n = 12) using a repeated-measures ANOVA. We observed a main effect of SETUP on comfort ratings (Greenhouse-Geisser corrected: $F_{1.3,14.6} = 4.53$, $p = 0.042$, $\eta p^2 = 0.29$). Descriptively, participants reported the highest comfort with the 64ch setup (mean ± SD: 1.50 ± 0.67), followed by Sushi (1.83 ± 1.03), and the lowest comfort with Surface (2.50 ± 1.31). Holm-corrected post hoc comparisons were not significant (all p ≥ 0.061), with the largest descriptive difference observed between Surface and 64ch (mean difference = 1.0).
